# Higher prevalence of homologous recombination-deficiency in lung squamous carcinoma from African Americans

**DOI:** 10.1101/651794

**Authors:** Sanju Sinha, Khadijah A. Mitchell, Adriana Zingone, Elise Bowman, Neelam Sinha, Alejandro A. Schäffer, Joo Sang Lee, Eytan Ruppin, Bríd M. Ryan

## Abstract

To improve our understanding of the longstanding disparities in incidence and mortality across multiple cancer types among minority populations, we performed a systematic comparative analysis of molecular features in tumors from African American (AA) and European American (EA) ancestry. Our pan-cancer analysis on the cancer genome atlas (TCGA) and a more focused analysis of genome-wide somatic copy number profiles integrated with tumor-normal RNA sequencing in a racially balanced cohort of 222 non-small cell lung cancers (NSCLC) reveals more aggressive genomic characteristics of AA tumors. In general, we find AA tumors exhibit higher genomic instability (GI), homologous recombination-deficiency (HRD) levels, and more aggressive molecular features such as chromothripsis across many cancer types, including lung squamous carcinoma (LUSC). GI and HRD levels are strongly correlated across AA tumors, indicating that HRD plays an important role in GI in these patients. The prevalence of germline HRD is higher in AA tumors, suggesting that the somatic differences observed have genetic ancestry origins. Finally, we identify AA-specific copy number-based arm, focal and gene level recurrent features in lung cancer, including a higher frequency of *PTEN* deletion and *KRAS* amplification and a lower frequency of *CDKN2A* deletion. These results highlight the importance of including minority and under-represented populations in genomics research and may have therapeutic implications.

## Introduction

In the United States, African Americans (AAs) have the highest cancer incidence and lowest survival across multiple cancer types ^1^. The reasons for these persistent trends are not clear. Our current understanding of the molecular mechanisms of tumorigenesis has primarily come from analyses of tumors derived from populations of European ancestry, including those based on TCGA ^2^ where only 8.5% of samples are from AAs. This raises a question about whether there is a difference in tumor evolution and molecular features by genetic ancestry. Recently, Yuan *et al.* compared somatic copy number alteration (SCNA)-based genomic instability (GI) across genetic ancestry in TCGA and found that breast invasive carcinoma (BRCA), head and neck squamous cell carcinoma (HNSC), and uterine corpus endometrial carcinoma (UCEC) tumors from AAs had significantly increased GI compared with tumors from European Americans (EAs) ^2^. Further, recent work has built a new consensus reference genome from DNA samples from individuals of African descent and found that the African pan-genome contains numerous large insertions—whose total length comprises ∼10% of the genome—that are not present in the current human reference genome (GRCh38) ^3^, which was primarily derived from a single individual of primarily European descent ^4^. Together these data highlight the need for studies specifically investigating the molecular landscape of cancer in minority and under-represented populations.

Population-specific molecular patterns in tumor biology and cancer genomics have been reported in recent years ^5-8^ with limited power and coverage. Here, we identified ancestry-specific molecular features in tumors across all cancer types for EA and AA in TCGA and with a special focus on lung cancer, which is the leading cause of cancer-related death and for which there are persistent disparities in both incidence and mortality ^9^. Our analysis confirms the higher genomic instability (GI) and also a higher prevalence of homologous-recombination deficiency (HRD) in tumors from AAs compared with pan-cancer and LUSC from EAs. This suggests a genetic ancestry-associated disparity in HRD, which we confirmed by finding a higher prevalence of germline HRD in tumors of AAs compared with EAs, again in both pan-cancer and LUSC. Further, we have performed a genome-wide copy-number features comparison in EAs and AAs to identify ancestry-specific arm, focal and gene-level features. Our results highlight the importance of including minority and under-represented populations in cancer genomics research and may have therapeutic implications.

## Results

### African Americans have higher GI and HRD

We mined TCGA somatic copy number profiles of 6,492 tumor samples originating from 23 different tumor types (Figure S1, Table S1) and observed significantly higher GI burden in AAs scaled within cancer types (Wilcoxon rank-sum (denoted *Wilcoxon* from here onwards) *P*< 6.9E-07, Figure 1A-top panel), consistent with previous observations ^2^. At the cancer type level, these differences were most significant in breast (BRCA), head and neck (HNSC), stomach adenocarcinoma (STAD), and cervical squamous cell carcinoma and endocervical adenocarcinoma (CESC) cancers, with a general trend towards higher GI burden in 17 out of the 23 cancer types (Figure S1). We repeated this analysis separately for CNV loss and gain based-GI and observed a consistent pattern (Figure S2, Supplementary notes 1).

**Figure 1:**
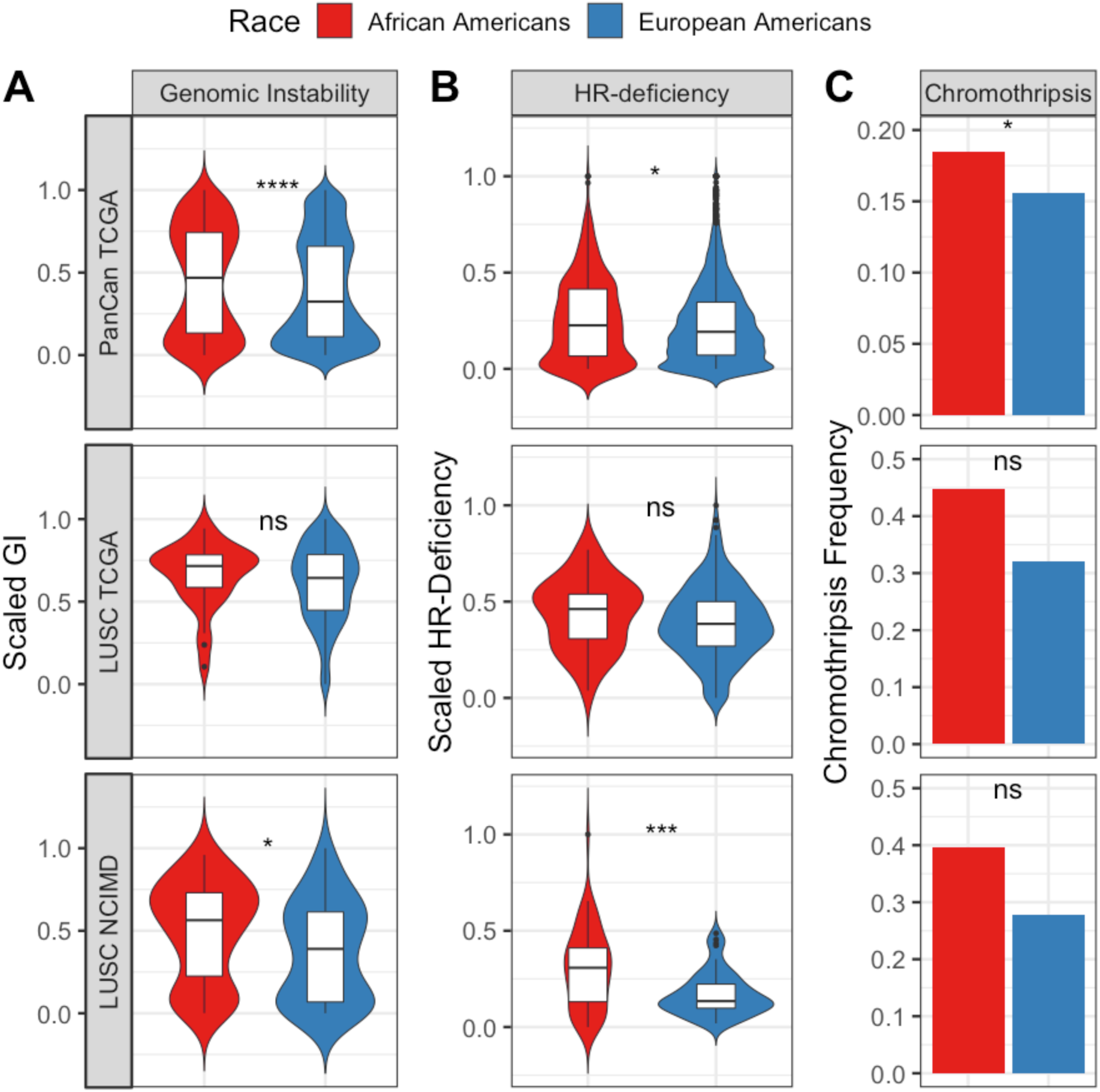
Differences in GI, HRD and chromothripsis across AA and EA genetic ancestry in pan-cancer scaled within each cancer-type and lung cancer. (A) Genomic instability, (B) homologous recombination deficiency and (C) chromothripsis quantified in pan-cancer TCGA samples (top most panel), LUSC TCGA samples (middle panel) and LUSC samples from NCI-MD cohort (bottom panel). Here, *ns* stands for non-significant, where *, ***, **** denotes <0.05, <0.001, < 0.0001.

We tested the hypothesis that higher GI across tumors from AAs is due to a higher prevalence or extent of HRD, which was previously identified as a key contributor to GI in cancer ^10^. Using the sum of loss of heterozygosity (LOH) events—a hallmark of HRD ^11,12^—as a surrogate, we observed a strong correlation between GI and HRD in both pan-cancer (Spearman Rho=0.56, *P* <2.2E-16) and cancer type-specific analyses (Figure S3A, Table S1), where, in AA tumors, the correlation observed is stronger than in EA tumors for both pan-cancer (Spearman Rho for AA=0.57, for EA=0.48, *P* <2.2E-16 for both) and cancer type-specific analyses (Table S2). We further observed that HRD scaled within cancer types is significantly higher in AAs in a pancancer (Figure 1B-top panel, Wilcoxon *P*< 1.5E-02). To seek additional support for these results, we quantified HRD using two additional signatures such as telomere allelic imbalance (AIL)—the number of regions of allelic imbalance that extend to one of the sub-telomeres but do not cross the centromere (Figure S3B)—and large-scale state transitions (LST)—the number of breakpoints between regions longer than 10 Mb after filtering out regions shorter than 3 Mb (Figure S3C) ^11^. These measures of HRD were significantly higher in AAs compared with EAs in a pan-cancer TCGA analysis using both individual measures as well as the net sum (Figure S4B-D, Wilcoxon *P*< 7.7E-02 for LST; *P*< 2.2E-02 for AIL; *P*<1.9E-02 for net sum). These additional two measures, as well as the sum of all three HRD measures, also correlate with GI (P <2E-16 for all, Spearman Rho = 0.47 for LST, 0.60 for AIL, 0.58 for net sum). Consistent with the pattern of GI and LOH-based HRD correlation across ancestry, these correlations are stronger in AA tumors compared to EA in both pan-cancer (AIL-based measure : Spearman Rho for AA=0.66, for EA=0.60, *P* <2.2E-16 for both; LST-based measure : Spearman Rho for AA=0.51, for EA=0.47, *P*<2.2E-16 for both) and cancer type specific analyses (Table S2). This suggests that HRD contributes to the ancestry-specific pattern of higher GI burden in AAs across cancer types. When analyzed by specific cancer type for the three HRD scores, we find that BRCA and HNSC had significantly higher HRD in AAs compared with EAs. A trend towards increased HRD among AAs was observed in 11 out of 17 cancer types where increased GI was also observed (Figure S3). We further confirmed these results by quantifying the contribution of mutational signature (*mutSig*) 3, known to be strongly associated with HRD ^13-15^, and found it to be higher in tumors from AAs in pan-cancer (*Wilcoxon* P<1E-03, Figure S4E). Testing each cancer type specifically for a higher *mutSig* 3 in AAs, we found that BRCA and HNSC have a higher prevalence of this HRD-related signature, which is consistent with the SNCA hallmarks-based quantification of HRD above (*Wilcoxon, P*<0.01 and <0.10, respectively, Figure S3E).

Moving from the pan cancer analysis to a focused study of lung cancer, both GI and four out of five measures of HRD mentioned above were higher in tumors from AAs compared with EAs in lung squamous carcinoma (LUSC), but the differences did not reach statistical significance (Figure 1A & 1B – middle panel). We reasoned that this could be due to the limited number of AA tumor samples (29 AA compared to 346 EA) and is supported by a power analysis of TCGA samples across ancestry (Supplementary notes 2). Thus, to test the hypothesis that GI and HRD is higher in lung tumors from AAs, we generated a genome-wide copy number profile of 222 non-small cell lung cancer tumor samples (cohort name – NCI-MD), 105 LUAD (AA=63, EA=42) and 117 LUSC (AA=63, EA=54) (Table S3) using the OncoScan platform, which provides a comprehensive coverage of 50-100 Kb in cancer genes and 300Kb in the rest of the genome (methods). Confirming our hypothesis, we found that LUSC tumors from AA had significantly higher GI compared with EA (Figure 1A-bottom panel; *Wilcoxon P*<6E-03). In contrast, in lung adenocarcinoma (LUAD), where comparative to LUSC more AA samples are present (52 AA, 387 EA in TCGA), both in TCGA and NCI-MD, there were no statistically significant differences in GI by ancestry (Figure S5A).

When HRD was quantified using the number of LOH events as a surrogate, we again observed a strong positive correlation between GI and HRD (Spearman Rho=0.64, P<2E-16) and also observed significantly higher HRD in AAs with LUSC (FDR adjusted *P*-value<2.0E-04, Figure 1B-bottom Panel), but not LUAD, which is consistent with our findings above (Figure S5B). Additionally, we performed multivariate linear regression to separately model GI and HRD in the NCI-MD cohort stratified by histology as a function of ancestry to account for potential confounding factors such as tumor stage, patient age, sex, smoking status and pack-years of cigarettes. Here, we found ancestry to be strongly associated in the consistent to before direction i.e. higher in AA with both GI and HRD (*FDR*<3E-02 and *FDR* <5.35E-05, respectively) only in LUSC and not LUAD (FDR< 0.24 and FDR<0.09, respectively), consistent with our previous observations (Table S4). When repeated for TCGA samples, we again observed a strong association of in the right direction in pan-cancer for GI (FDR<2.2E-07) and for all different measures of HRD (FDR <4.5E-06 for LOH, <7.8E-07 for AIL, <3.8E-05 for LST, <2E-01 for *mutsig* 3, Table S4). This concludes that after adjusting for tumor stage, gender and smoking etc, the observed ancestry differences remain consistent in both NCI-MD and TCGA.

In NCI-MD, we infer the ancestry in an unsupervised manner via a PCA of all the SNPs (Figure S6, methods) and cluster via support vector classification (SVC). We found that inferred ancestry is concordant with self-reported ancestry for 98.3% of the cases, where four samples were misclustered and deviated from self-reported ancestry (Table S1). We removed these samples and repeated the whole analysis above and found completely consistent results with comparable significance (higher GI and HRD in LUSC AA *Wilcoxon P*<6E-03 and P<2E-04).

### AAs have more complex structural variants in pan-cancer

The observed deficiencies in DNA damage repair related with GI, especially in LUSC, prompted us to chart the landscape of structural variants recently reported to be related with HRD ^16^. We studied chromothripsis (CHTP), which was first described as a one-time catastrophic event that leads to chromosome shattering and tens to hundreds of simultaneously acquired oscillatory copy number aberrations clustered to one chromosome ^17,18^.

Using the classical definitions, i.e., many oscillatory (one type of CN) copy number events clustered on a chromosome (methods) ^17,18^, tumor samples with CHTP have significantly higher HRD than samples without CHTP (*Wilcoxon* P<3.2E-10, for all four HRD markers) in both pancancer (TCGA) and LUSC (NCI-MD), suggesting that the above proposed relationship between CHTP and HRD is present across pan-cancer. Consistent with the trends outlined above, we observed a higher frequency of CHTP in tumors from AAs compared with EAs in pan-cancer samples from TCGA (Figure 1C-top panel, Fisher’s one-sided test *P*<0.028, Odds Ratio (OR)= 1.25), in LUSC samples from TCGA (Figure 1C-middle panel, *P*<0.11, OR=1.4,), LUSC from NCI-MD (Figure 1C-bottom panel, *P*<0.12, OR=1.24) and also in LUAD from TCGA and NCI-MD (Figure S5C, S6) but to a weaker extent. These patterns were consistent when adjusted for age, sex and stage across both cohorts (multivariate regression P for ancestry <2.8E-03) and further when a recently described CHTP definition – where two oscillation state were allowed in the CHTP affected region was used (methods, Table S1). We further analyzed the chromosome frequency distribution of CHTP, which varied by histological subtype and ancestry (Figure S7B). Similar ancestry-specific patterns across TCGA and NCI-MD in chromosome enrichment of CHTP were observed on chromosomes 2 and 12 among AAs compared with EAs in LUSC (Figure S7B-C).

### Higher germline prevalence of HRD in African Americans

Given the higher prevalence of HRD in AA tumors across pan-cancer and LUSC, we mined germline pathogenic variants across 10,389 adult-tumors in TCGA (methods described here ^19^), and investigated whether cancer patients of AA descent are more likely to have a predisposition for HRD than EAs by testing whether AAs are more likely to have germline pathogenic variants related to HRD ^19^. In TCGA, both pan-cancer and LUSC, but not LUAD, we found AAs were significantly more likely to have germline HR-deficiency, where at least one of the genes in the HR-pathway ^10^ has a pathogenic variant. (Figure 2-left panel, OR=1.2; *P*< 0.02 for pan-cancer; Figure 2-right panel, OR=6; *P*< 8E-04 for LUSC; Figure S8, *P*<0.22 for LUAD).

**Figure 2:**
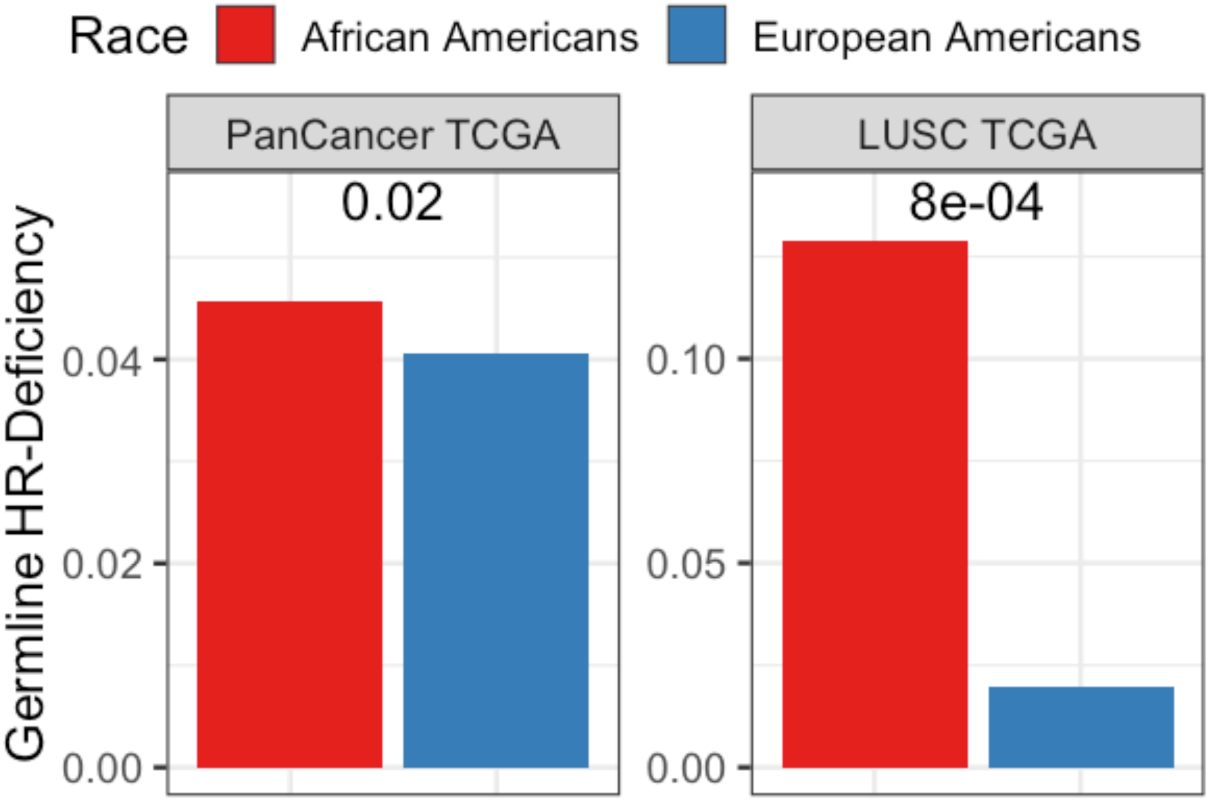
Germline-HRD across ancestry: Prevalence of germline HRD in AA and EA calculated using total frequency of germline pathogenic variants in HR-pathways genes, in pancancer (left) and LUSC (right) from TCGA where the significance was calculated using a Fisher’s test.

### The landscape of arm- and focal-level SNCA in AA and EA lung cancer

To identify SCNA-based ancestry specific features in detail, we examined population-specific patterns in lung cancer for chromosome arm- and focal-level (shorter than half a chromosome arm) events in the NCI-MD study where statistical power for both populations was available (Table S3). Further support for key observations was demonstrated in TCGA. Recurrent arm-level and focal-level events were identified for both populations separately using GISTIC ^20^ (methods, FDR<0.1) and used to map genome-wide copy number map across histology and ancestry (Figure 4, Tables S5-6).

For each chromosome arm, alteration frequency and recurrence significance by ancestry for both aberrance type—amplification and deletion—was charted for NCI-MD and TCGA (Figure 3).We identified known cancer-specific arm-level events, including amplification of 3q and 5p and deletion of 3p ^21^, in both populations (Table S5). Similarly, 19p deletion, a molecular signature of LUAD ^8^, is recurrent in EAs and AAs at a similar frequency of ∼45% (Figure 3, Table S6). Recurrent population differences were observed with unknown relevance; for example – LUSC arm level deletions on chromosome 4p and 4q and LUAD amplifications of 7p and 7q occurred at higher frequency in AAs compared with EAs and was replicated in TCGA (Figure 3, Table S5-6).

**Figure 3:**
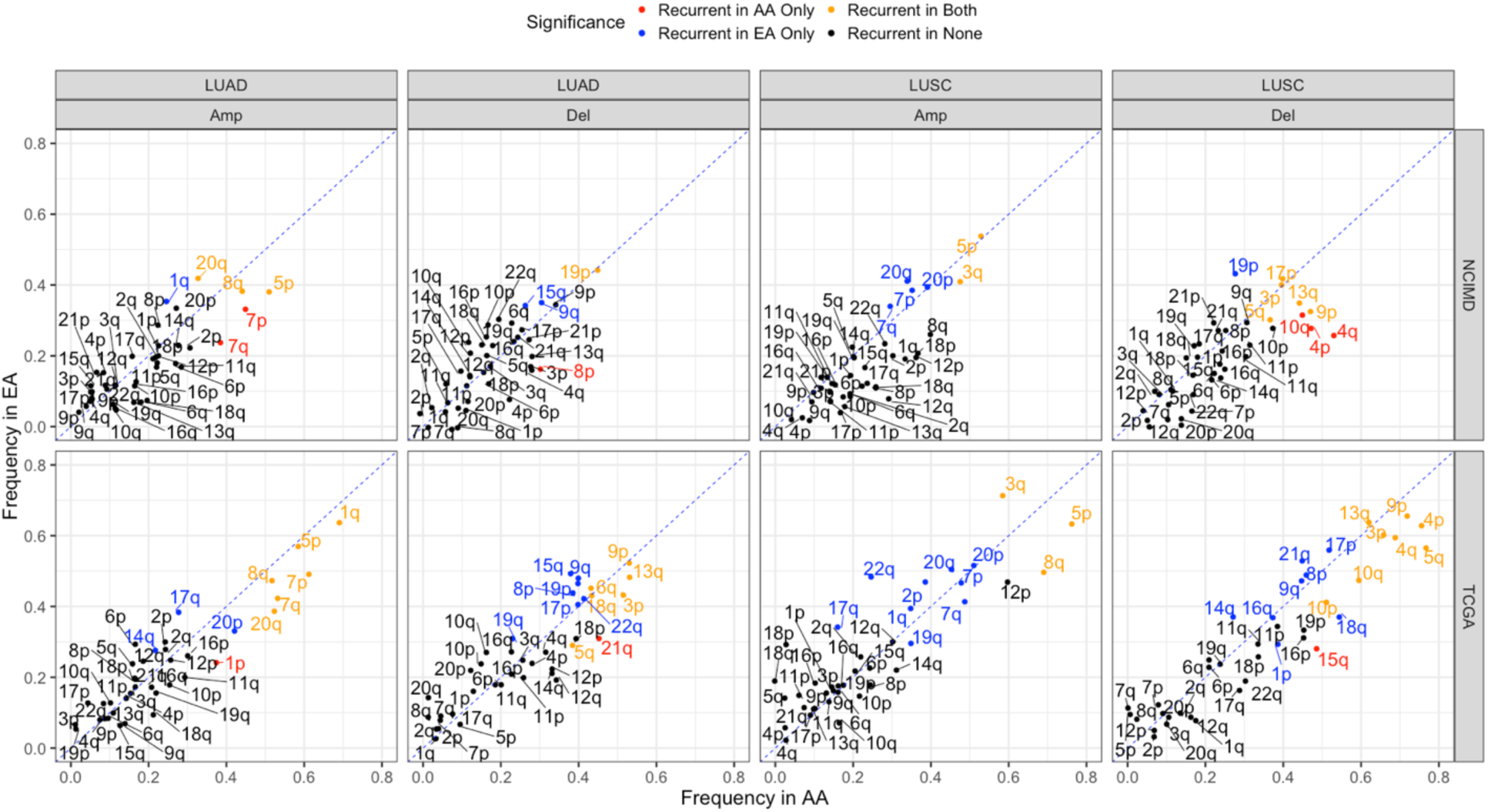
Characterization of arm-level events in lung cancer across populations. Frequency distribution of aberrant events on autosomal chromosome arms (Del-deletion and Amp-amplification) in LUAD and LUSC for NCI-MD and TCGA. The diagonal dashed line denotes the null line with points falling away from this line indicating chromosome arms with alteration frequency differences across populations. A color code is provided for population-specific recurrence.

To visualize genome-wide focal-level SCNA events across population, including co-occurrence and mutual-exclusivity, we created a global SCNA map showing genome-wide SCNA frequency distributions for both LUSC and LUAD (Figure 3-left panel, Tables S7-8). The overlap in recurrent focal regions among EAs and AAs was 59% and 70% for LUAD and LUSC, respectively (Figure 4-left panel). Further, we observed population-specific patterns of cooccurring and mutually exclusive SNCA events (Figure 4-left panel). To identify potential novel AA-specific copy number driven focal-level regions, we filtered and selected high-confidence recurrent focal-level regions from GISTIC whose 1) alteration frequency is greater than 5% in AAs, 2) frequency was at least two times higher in AAs than EAs and 3) were recurrent only in AAs and making sure that no recurrent peak of the same type (amp or del) is present in EAs within the region or an extended additional 10% on both sides of the region length. We identified eight such potential AA-specific potential driver regions. The top hit ranked by significance is 22q11.23 in LUSC (Figure 4-right panel) with a frequency of 27% in AA and 13% in EAs. Based on previous studies ^22^, we tested whether this event is somatic or germline by investigating matched normal tissue samples and observed that 2/5 normal samples from AAs also have deletion of 22q11.23, suggesting that this event could be germline (Table S9). Notably, this frequently deleted region in LUSC on 22q11.23 appears to be disjoint from the full extent of the nearby region on 22q11.21 that is hemizygously deleted in DiGeorge syndrome. The region with the second highest fold change in alteration frequency in LUSC – 12p12.1 (Figure 4-right panel), is a short length region comprising *KRAS* and is discussed in detail in next section. Thirdly, common to both LUAD and LUSC, the 20p12.1 region is more four times more deleted in AAs. This region includes the genes *FLTR3* and *MACROD2*.

We also identified several SCNA events previously linked with AA ancestry in cancer. We replicated an AA-specific amplification of the oncogene *KAT6A* in LUAD ^*23*^ and also identified a recurrent deletion of 4q35.2 extending to telomeres in LUSC, which has been recurrently found in co-morbid psychiatric illness ^46^ and comprises the gene *FBXW7* previously shown to be deleted in colorectal cancer and triple negative breast cancer among AAs ^24,25^ (Table S10). Relevance of both of these observations are yet unknown. In LUAD, two close regions on 8q24 (8q24.3 and 8q24.21) were significantly amplified in AAs only (frequency =33% and 18% in AAs and EAs, respectively). Within 8q24.3, *PVT1* CN profile was significantly associated with expression (*P*<7E-03) while *HSF1, DGAT1* and *BOP1* were also significantly associated with expression (*P*<7E-03) (Table S11).

**Figure 4:**
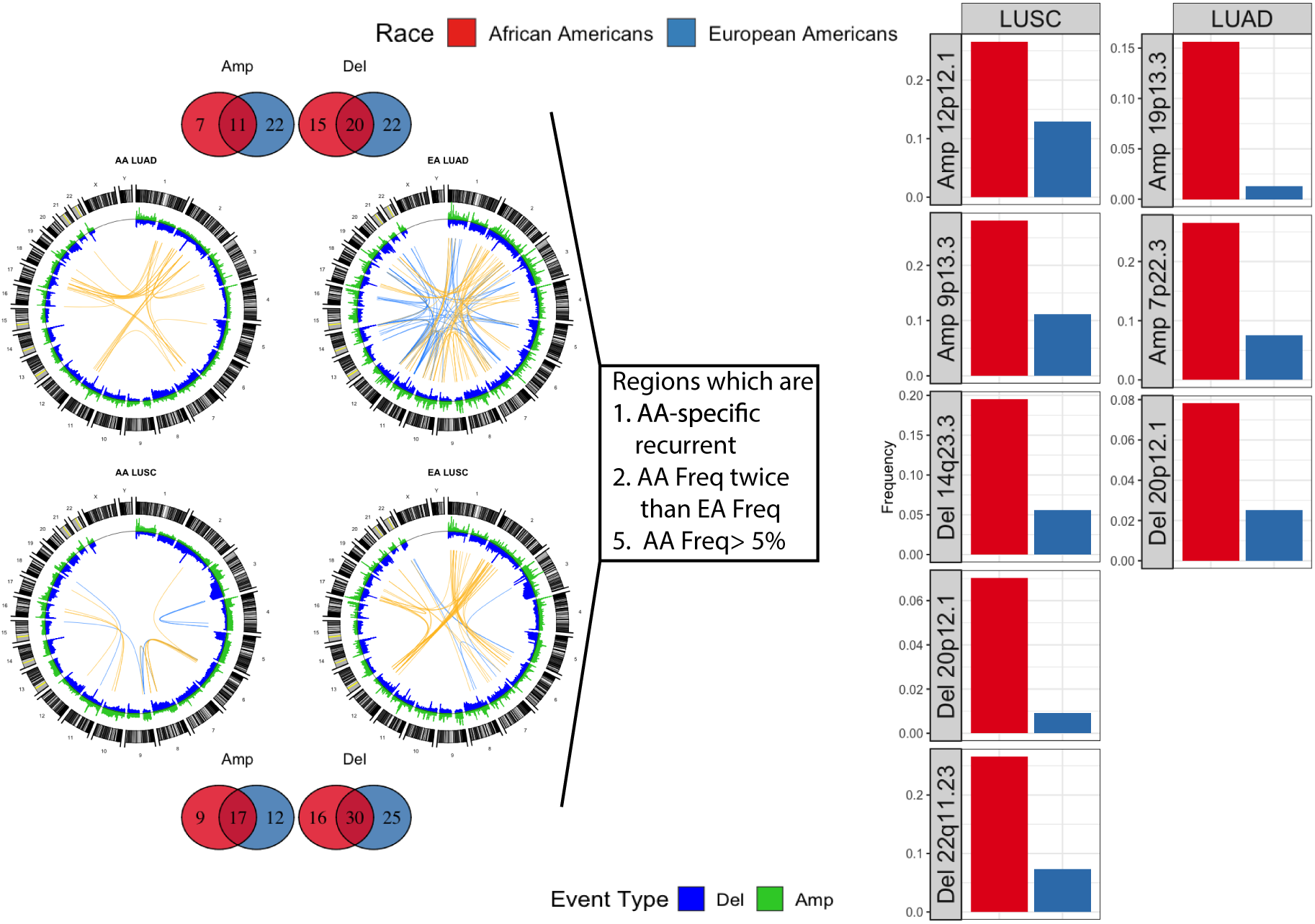
Global SNCA map in lung cancer across population (AA and EA) and histology (LUSC and LUAD). Segment deletions and amplifications are blue and green, respectively in the left panel Circos plot. Number of overlap and unique recurrent regions between AA and EA in LUSC and LUAD are shown via Venn diagrams at the top and bottom. Highly positively (cooccurring) and negatively (mutually-exclusive) correlated CN segments are connected with yellow and blue arcs, respectively. The top 50 segment pairs are chosen with Pearson-Rho>0.75 for positive correlations and Pearson-Rho < −0.50 for negative correlations between SCNA events. Steps provided in the central box are used to identify CNV-driven AA-specific potential-driver regions whose cytoband and frequency in AA and EA are provided in right via bar plots in the respective subtype of lung cancer.

### Landscape of driver genes in AA and EA lung cancer

We systematically analyzed the recurrence and alteration frequency of known lung cancer tier-1 driver genes mined from the cancer gene census of COSMIC (Figure 5A). We identified population-specific SCNA patterns of drivers significantly correlated with gene expression (Figure 5A and Figure S9-10). In LUSC, *KRAS* and *PTEN* are recurrently amplified and deleted, respectively, in AAs only (*KRAS* amp frequency:23% in EA compared to 51% in AA). *CDKN2A* is recurrently deleted in both populations, but the frequency is 64% in AAs compared with 35% in EAs (Figure 5A & 5B). These three population-specific patterns are consistent and supported by TCGA (Figure S9-10).

**Figure 5:**
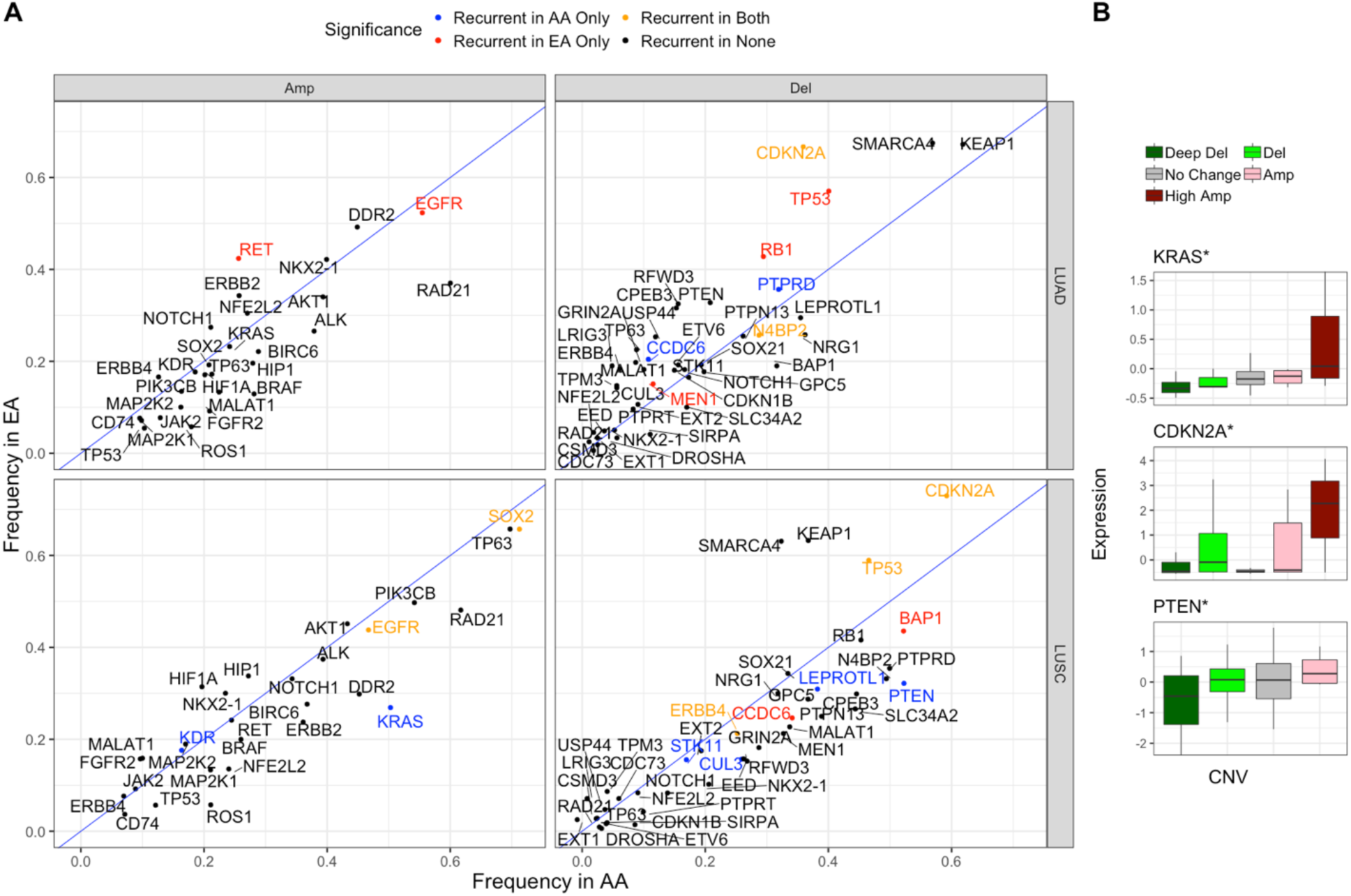
Lung cancer drivers across population. **(A)** Driver-genes amplifications and deletion frequency of lung cancer driver genes across population and histology. **(B)** Effect of copy number change on expression of drivers with population-specific patterns.

## Discussion

Here, we mapped molecular features of EA and AA tumors across many cancer types and further, with greater depth and power, in lung cancer. We observed that, consistent with previous reports ^2^, GI is higher in AAs across multiple cancer types. The higher GI is unlikely to be related to the recently identified unmapped 10% of the genome that is found in populations of African ancestry^3^ as we found both copy number gain- and copy number loss-based GI to be higher in AAs. We hypothesize and confirm that this higher GI is likely due to a higher prevalence of HRD in tumors from AAs. Further supporting this, we also identified a significantly higher prevalence of mutational signature 3—closely associated with HRD^13-15^— among tumors from AAs. We further show that tumors from AAs have a higher frequency of aggressive molecular features including structural variants (chromothripsis).

Higher SNCA-based GI and HRD in tumors from AAs raises the question of whether underlying defective DNA repair mechanisms could drive this observation. While HRD has been linked with germline and somatic mutations in *BRCA1/2* ^26^, no striking differences in the somatic mutation frequency of these genes has been demonstrated in cancer between EAs and AAs ^2,27^. To investigate whether the increased HRD could be driven by a germline event, we leveraged the recent identification of germline pathogenic variants ^19^ and identified a higher proportion of HRD-related pathogenic variants among AAs compared with EAs, suggesting that GI/HRD events and tumor evolution could be shaped by these features. The observation that several cancer types occur at an earlier age among AAs ^28^ and evidence that germline pathogenic events are associated with early-onset disease ^19^ are consistent with these data.

Higher HRD in LUSC suggests that these tumors could better respond to PARP inhibitors and that perhaps, AAs may be more likely to respond. PARP inhibitors are not commonly used in non-small cell lung cancer treatment, though in combination with chemotherapy they have shown promising efficacy in both cell lines ^29^ and a clinical trial ^30^. Of note, in the latter study, the benefit from the combination treatment was primarily restricted to LUSC tumors. In the future, clinical studies focusing on an enrichment of treatment response based on HRD features and potential population differences should be conducted.

When we looked in granular detail at SCNA events in LUAD and LUSC, we found several regions to be recurrently altered only in AAs and several regions previously linked to disparities in other cancer types, including 4q35.2, where *FBXW7* is also associated with expression (of note, arm level events on 4q were also recurrently only among AAs). This gene controls proteasome-mediated degradation of oncoproteins including cyclin E, c-Myc, mTOR and Jun ^31^. It has also been implicated in DNA repair ^32^. Interestingly, recurrent deletions of *FBXW7* have been associated with metastatic ^24^ and is deleted in triple negative breast cancer ^25,33^ and colorectal cancers in AAs ^7^. Amplification on 8q24.3 and 8q24.21 also occurred in AAs at a higher and significant frequency in AAs than in EAs. In this region, *BOP1, DGAT1* and *HSF1* and *PVT1* were associated with expression. Of note, we also validated the previous observation ^34^ that the oncogene *KAT6A* ^*23*^ is amplified among AAs only (18% vs. 0%) and further find that this locus is associated with LUAD and gene expression. Inhibitors of this histone acetyltransferase have been developed and induce senescence ^23^, raising the possibility that they could be useful in *KAT6A*-driven cancers. As our study used the same platform to compare SCNA events across EAs and AAs, it largely removes the possibility that technical artifact could confound our observations. Also, the dataset is well balanced across EA and AA for both LUAD and LUSC.

In summary, in a pan-cancer analysis we have identified population differences in molecular features, including GI and HRD. As these features are related to therapy response ^11,35,36^, these findings could have therapeutic potential. We also find higher GI and HRD in LUSC and highlight some granular differences at the SNCA level in canonical lung cancer genes, such as *CDKN2A, KRAS* and *PTEN*. Further, defining these differences in both genome-wide and more focal regions gives rise to distinct differences in lung tumor biology between AAs and EAs and supports recent work showing that inherited variants and thereby, genetic ancestry, can shape tumor grade evolution at a molecular level and influence the somatic nature of a tumor ^37^. These results may provide insights on therapy decisions and have implications for future therapeutic strategies and/or the recruitment of patients into clinical trials. Finally, our work highlights the importance of including under-represented populations in balanced genomic studies to study population-specific mechanisms underlying molecular patterns and cancer evolution.

## Methods

### A. Samples preparation and processing

#### Sample characteristics

Patients living in the Baltimore Metropolitan area with histologically confirmed cases of lung adenocarcinoma (LUAD) and lung squamous cell carcinoma (LUSC) were prospectively recruited to the ongoing NCI-MD Case Control Study ^38^. Institutional Review Boards at seven participating Baltimore hospitals and the NCI approved the study with written informed consent obtained from all patients. We conducted a retrospective study of eligible participants that self-reported as AA or EA, with non-Hispanic ethnicity. Additional clinical and sociodemographic data for each patient were obtained from medical records and pathology reports. Macro-dissected primary lung tumor tissues were obtained from patients directly after surgical removal. Samples were placed in collection tubes, flash frozen and stored at −80°C until the OncoScan analyses were performed. Sample characteristics for the patients in which tumor DNAs were extracted from can be found in Table S1 (*n*=142 AA, 108 EA).

#### DNA extraction

DNA was extracted from fresh frozen micro-dissected primary lung tumor tissues using the Qiagen DNeasy Blood and Tissue kit spin column procedure according to the manufacturer’s protocol (Qiagen, Valencia, CA). Isolated primary lung tumor DNAs were initially quantified using a DS-11 spectrophotometer (DeNovix, Wilmington, DE). Subsequent Qubit fluorometer analyses were performed to assess DNA integrity and ensure the presence of intact double-stranded DNA of all samples (Invitrogen, Carlsbad, CA). DNA with a A260/A280 ratio between 1.8 and 2.0, a minimum concentration of 12 ng/µL, and a total concentration of 80 ng was used for further analysis.

### B. Preprocessing of Raw files

#### Genome-wide copy number analysis and data QC

DNA samples were sent for genome-wide copy number analysis using the Affymetrix OncoScan CNV array and run according to suggested manufacturer protocols. The OncoScan array is based on molecular inversion probe technology and provides comprehensive high-resolution copy number detection across the genome and at pan-cancer driver genes. OncoScan fluorescence array intensity (CEL) files were converted to OSCHP files using the hg19 reference (OncoScan_CNV.Ref103.na33.r1.REF_MODEL reference file included with the Affymetrix OncoScan Console software, version 1.3). Manual re-centering of samples was performed by adjusting the TuScan log_2_ R using the OncoScan Console. Clonality analysis was performed with Affymetrix OncoClone Composition tool.

#### Segmentation of NCI-MD and TCGA intensity files

For these samples, the Chromosome Suite Analysis (CHAS) was used for segmentation of intensity files at the default hyperparameters for output of segments with their copy number, log_2_R and B-Allele frequency (BAF) information. For TCGA samples, Level 3 – segmented files were retrieved from via firehose pipeline where consistent version of reference (hg19) was used.

#### Purity and Ploidy calculation

Using OncoClone tool, provided by Affymetrix which uses the algorithm ASCAT ^39^, we have computed the purity and ploidy of samples from our cohort (Table S16 and Supplementary Note 3). Further, intratumor heterogeneity (ITH) was calculated using TuScan^28^ algorithm, a further extension of OncoClone.

### C. Quantifying GI, HRD and Chromothripsis

#### Quantification of genomic instability

For every sample in TCGA and NCI-MD, genomic instability was defined by ratio of the total length of regions with copy number other than 2 to a constant of 3.3E9 based on previous works ^40,41^. For TCGA samples, we only selected cancer types with at least five AA samples.

#### Homologous-Recombination deficiency (HRD) quantification based on Mutation Signature

Mutation signature 3 was mined from mSignatureDB ^42^, a database of mutation signatures for more than 15,000 tumor samples from more than 73 projects.

#### Quantifying germline HRD using germline pathogenic variants for HR-pathway genes

We charted the status of HRD at a germline level for each patient in TCGA cohort by mining the pathogenic variants callings from the sequencing of normal blood or matched tissue samples from this work^19^. All the variant calls were downloaded from the database of Genotypes and Phenotypes (dbGaP).

#### Quantification of chromothripsis

With an aim to identify whether an autosomal chromosome has undergone through chromothripsis using CNV profile, we used four copy number based-hallmark traits of regions undergone chromothripsis. Some of these hallmarks of chromothripsis has undergone an evolution since its first description, hence we used two partially overlapping settings of hallmarks to identify chromothripsis based on the conventional ^43^ one and the most recent ^44,45^ description provided below. Chromosomes having all the following four hallmark properties are considered to undergone chromothripsis.

We modelled the four hallmarks chromothripsis^17^ via two tests for each sample. First, we filtered for chromosomes with significantly more events than the sample’s background, derived from all other autosomes. Specifically, a chromosome must have a higher number of copy number events than the median number of copy number events per chromosome in the sample. Second, for every chromosome that passed the first test, the distance between the events breakpoints on the chromosomes should be significantly lower than the background distribution of copy number events breakpoints within the rest of the chromosomes. To this end, we tested whether the distances between the breakpoints of events of given a chromosome was lower than the background distribution of distances between the breakpoints of events on the rest of the chromosomes. If not, we removed the terminal event with higher breakpoint distance from the penultimate and repeated the above step.

The above iteration was repeated for a chromosome until we found a region having greater than five events with significantly lower breakpoint distance (Clustered, FDR corrected *P*< 0.1) and the region comprise only one type of copy number events (Oscillatory CN State) at maximum. We repeated the above steps with a single modification to model and detect CHTP based on the recent definition^18^, where in a CHTP regions two oscillatory CN states or two types of copy number events can be there.

#### Association of copy number change with expression

For this study, total-RNA sequencing was performed for 56 out of 222 samples with CNV profile (31 LUAD & 25 LUSC). The association of copy number with expression was calculated via a one-tailed Wilcoxon rank-sum test, where samples were divided into two categories by thresholding on the median gene copy number to test, in a genome-wide fashion for each gene, whether samples with copy number higher than the gene median copy number in the cohort has expression significantly higher than the rest of the samples.

### D. Focal and arm events by GISTIC

#### Generating a copy number map with focal-, arm-level events via GISTIC

The GISTIC algorithm was used to find recurrent regions from the segmented file generated from CHAS. We used the following hyperparameter configuration throughout the study to find recurrent regions “*-genegistic 1 -smallmem 1 -broad 1 -brlen 0.5 -conf 0.90 -armpeel 1 – savegene 1*”. Based on this configuration, a gene GISTIC algorithm was used where *arm* level events are defined as aberrant regions with at least the length of half an arm and regions below this threshold are defined as *focal*. The confidence level used to calculate the region was 0.90 and the q-value was the default of 0.25.

#### E. Unsupervised ancestry inference via Principal component analysis (PCA)

Genotype information for 217611 SNPs were generated from OncoScan OSCHP file via apt-tools for the samples from NCI-MD. In this matrix, where each row represents a patient and each column represents a SNP, we performed a PCA with rank two, constraining the number of principal components (PC) to two (Figure S6). Next, we clustered the two PCs using support vector classification (SVC) with a linear kernel to identify two clusters. These two clusters were then tested for concordance with self-reported ancestry.

## Supporting information

supp_textandfigure

## Acknowledgements

We thank members of the Lab of Human Carcinogenesis, Curt Harris and Cancer Data Science Lab for insightful suggestions during the steps of our work. SS gratefully acknowledges the support of NCI-UMD Cancer Research Training Fellowship.

