## Supplementary material for "Higher prevalence of homologous recombination-deficiency in lung squamous carcinoma from African Americans": supp_textandfigure

**Supplementary Notes**

1. CNV-gain and -loss based genomic instability analysis
2. Statistical power analysis of TCGA samples from various populations
3. Qualitative characterization of NCI-MD cohort tumor samples
4. Arm aberration frequency negatively correlated with #genes present on the arm
5. Cancer genes in recurrent regions of LUAD and LUSC

**Supplementary Figures:**

**Figure S1:** Landscape of genomic instability (GI) across EAs and AAs in 23 cancer types in TCGA.

**Figure S2:** Gain and loss GI burden across EAs and AAs.

**Figure S3:** HRD across 23 cancer types in TCGA quantified via various scores.

**Figure S4:** HRD in pan-cancer in TCGA based on various hallmarks

**Figure S5:** GI, HRD and chromothripsis in LUAD from NCI-MD

**Figure S6:** Upsupervised ancestry inference by principal component analysis

**Figure S7:** Charting chromothripsis frequency across 23 cancer types in TCGA by population.

**Figure S8:** Charting prevalence of germline predisposition for HRD by population for 11 cancer types

**Figure S9:** Charting lung cancer drivers SNCA frequencies by population in LUAD and LUSC from TCGA.

**Figure S10:** Effect of copy number changes on expression for lung cancer drivers whose frequencies across populations are significantly different.

**Supplementary Notes**

**1.CNV-gain and -loss based genomic instability (GI) analysis**

***For TCGA Pan-Cancer***

We calculated CNV-gain and CNV-loss based GI and consistently observed both GI measures to be higher in AAs (Wilcoxon rank-sum $P$<5.2E-06 and $P$< 1.5E-06). Further, the trend of higher GI was observed in 16 out of 23 cancer types for both CNV-gain based and CNV-loss based GI (Figure S2A-B).

***For NCI-MD LUSC***

CNV-gain and CNV-loss based GI are calculated for LUSC from the NCI-MD cohort. We observed only CNV-loss based GI to be significantly higher in AAs (P < 4.5E-06 and <0.34, respectively).

**2. Statistical power analysis of TCGA samples from various populations**

We observed a negative correlation between the FDR-corrected significance for AA having higher GI and the proportion of AA samples included per cancer type, which was higher than expected when permuted a million times. (Spearman Rho=-0.34, $P$<0.15; empirical p<1E-04), suggesting that under-representation of samples from AAs is a limiting factor in terms of statistical power when comparing these two populations in TCGA.

**3. Qualitative characterization of NCI-MD cohort tumor samples**

Purity—the percentage of the tumor cell fraction within a sample—was successfully resolved in 194 out of 222 samples (Table S1). The mean purity was 34%. LUSC tumor samples had a significantly higher purity than LUAD, consistent with TCGA (mean purities in LUAD and

LUSC were 30.5 and 38.5, respectively; Wilcoxon rank-sum $P<0.009$) where an overall mean ploidy is 2.22.

**4. Arm-level aberration frequency negatively correlated with #genes present on the arm**

Broad level events across chromosome arms were quantified and plotted against the number of protein expressing genes [1] where a general trend of negative correlation between the frequency of an aberration on a chromosome arm and the number of genes present on the same arm (median *Spearman Rho*=0.41).

**5. Cancer genes in recurrent regions of LUAD and LUSC**

For both LUSC and LUAD, an analysis of known driver pan-cancer oncogenes and tumor suppressors mined from COSMIC [2] present in the recurrent regions indicated a limited overlap of cancer genes with these recurrent regions (Table S12) where we find that almost three-quarter of the deletion and amplified recurrent regions do not have any tumor suppressor or oncogene, respectively.

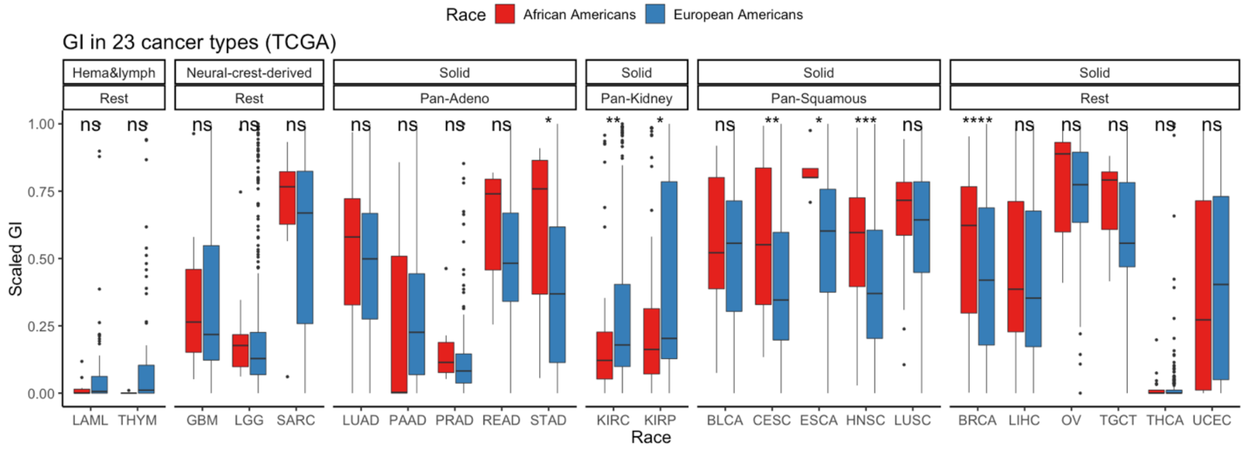


***Figure S1: Landscape of genomic instability (GI)*** *in AAs and EAs in 23 cancer types where at least five samples of AA are available in TCGA. Here, cancer types are clustered by cell type and tissue type of origin.*

**A.**

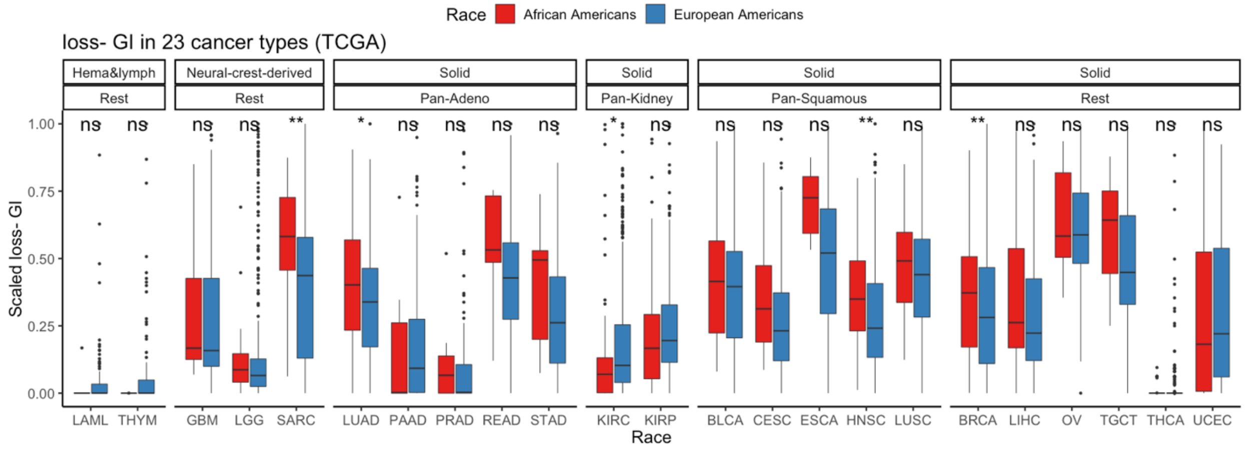


**B.**

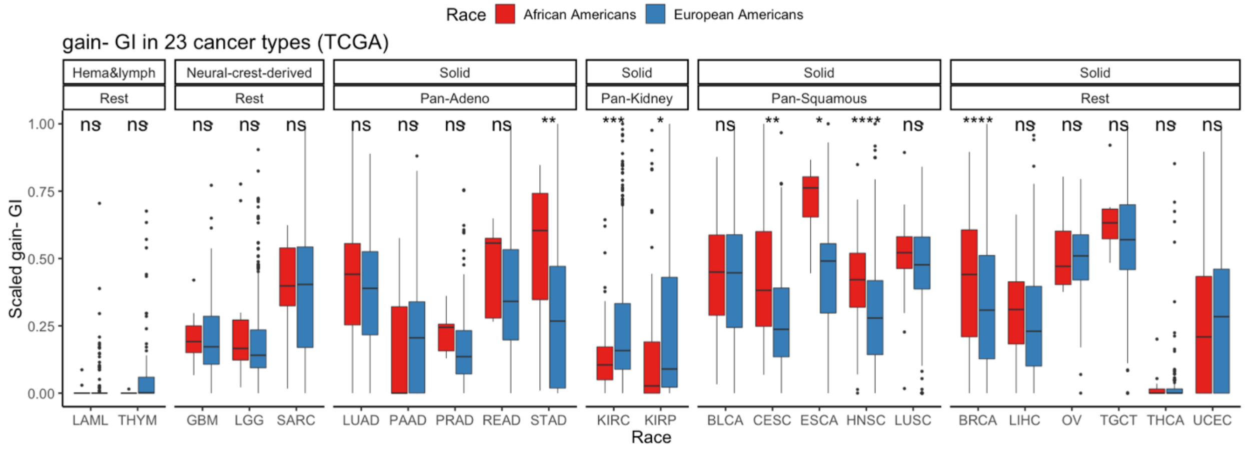


***Figure S2: Gain and Loss GI burden across populations.*** *A) CNV-gain and B) CNV-loss based genomic instability in AAs and EAs in 23 cancer types in TCGA.*

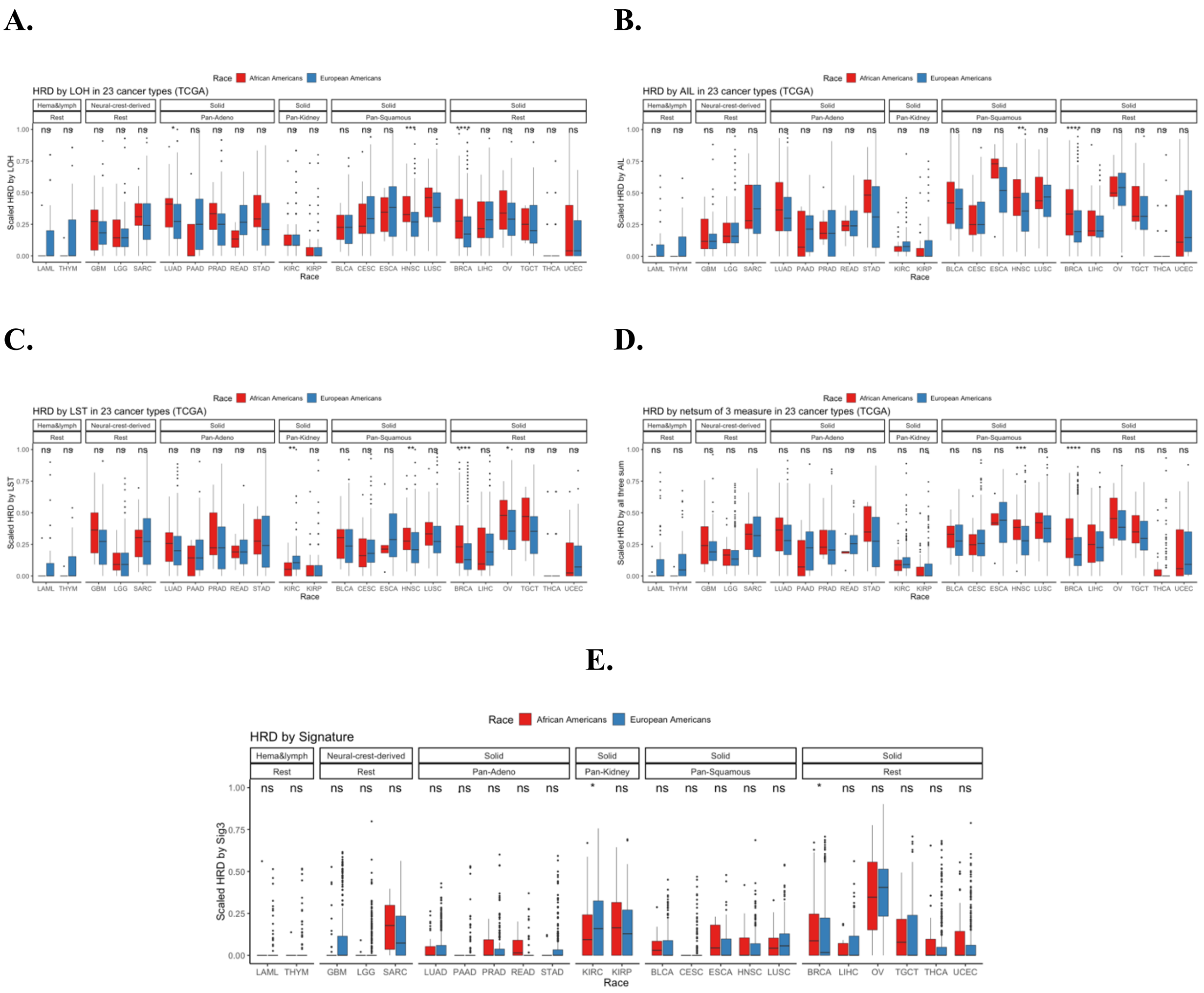


***Figure S3: HRD across 23 cancer types in TCGA quantified via score based on* (A*)*** *number of LOH events,* **(B)** *telomere allelic imbalance (AIL)* **(C)** *large-scale state transitions (LST)* **(D)** *Scaled Net sum of previous three defined as "genomic scar"[3]* **(E)** *Mutation signature 3 in AAs and EAs in various cancer types in TCGA.*

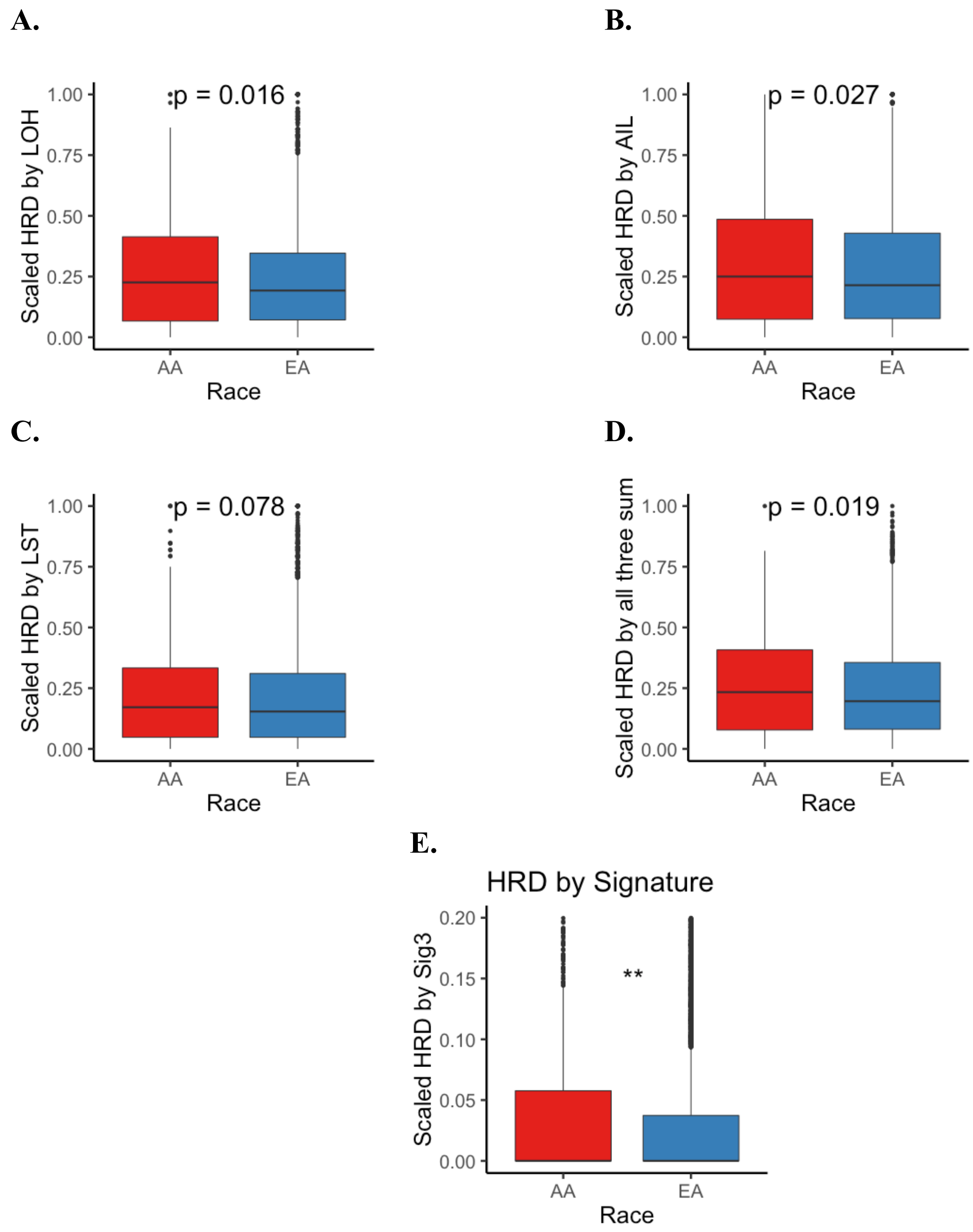


***Figure S4: HRD in pan-cancer in TCGA*** *quantified via score based on* **(A***) number of LOH events,* **(B)** *telomere allelic imbalance (AIL)* **(C)** *large-scale state transitions (LST)* **(D)** *sum of previous three defined as "genomic scar"***.**

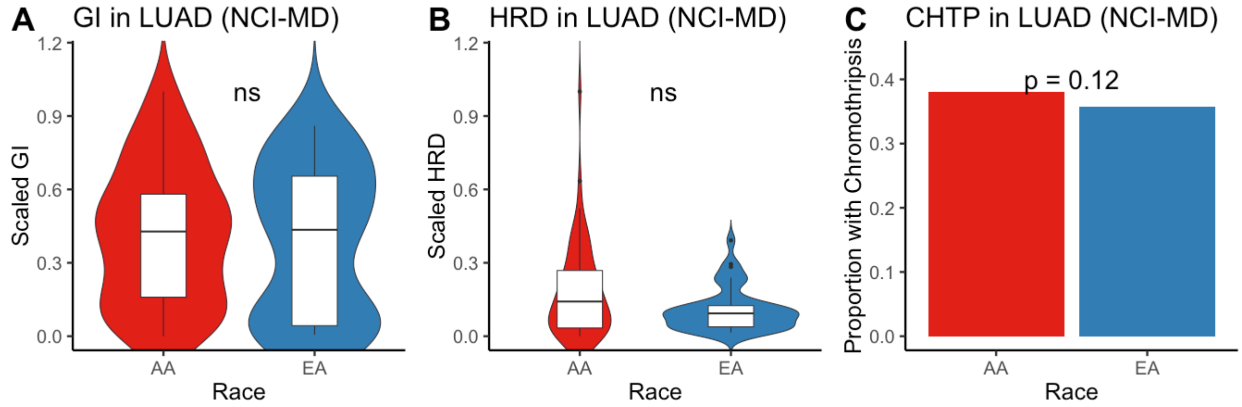


***Figure S5:*** *In LUAD from NCI-MD, we quantified* ***A)*** *GI and* ***B)*** *HRD via score based on number of LOH events* ***C)*** *Chromothripsis across AAs and EAs.*

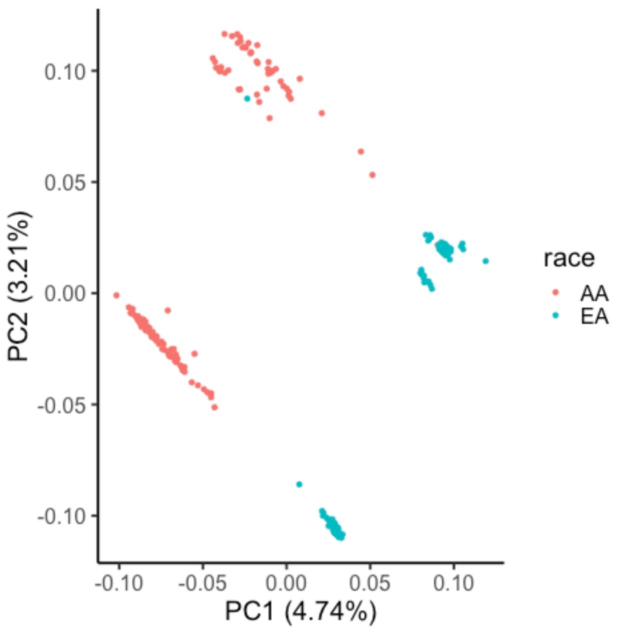


***Figure S6: Unsupervised inference of Ancestry.*** *Two major principal components (PCs) are provided colored by self-reported ancestry. These were used in unsupervised clustering via SVC to identify to two clusters.*

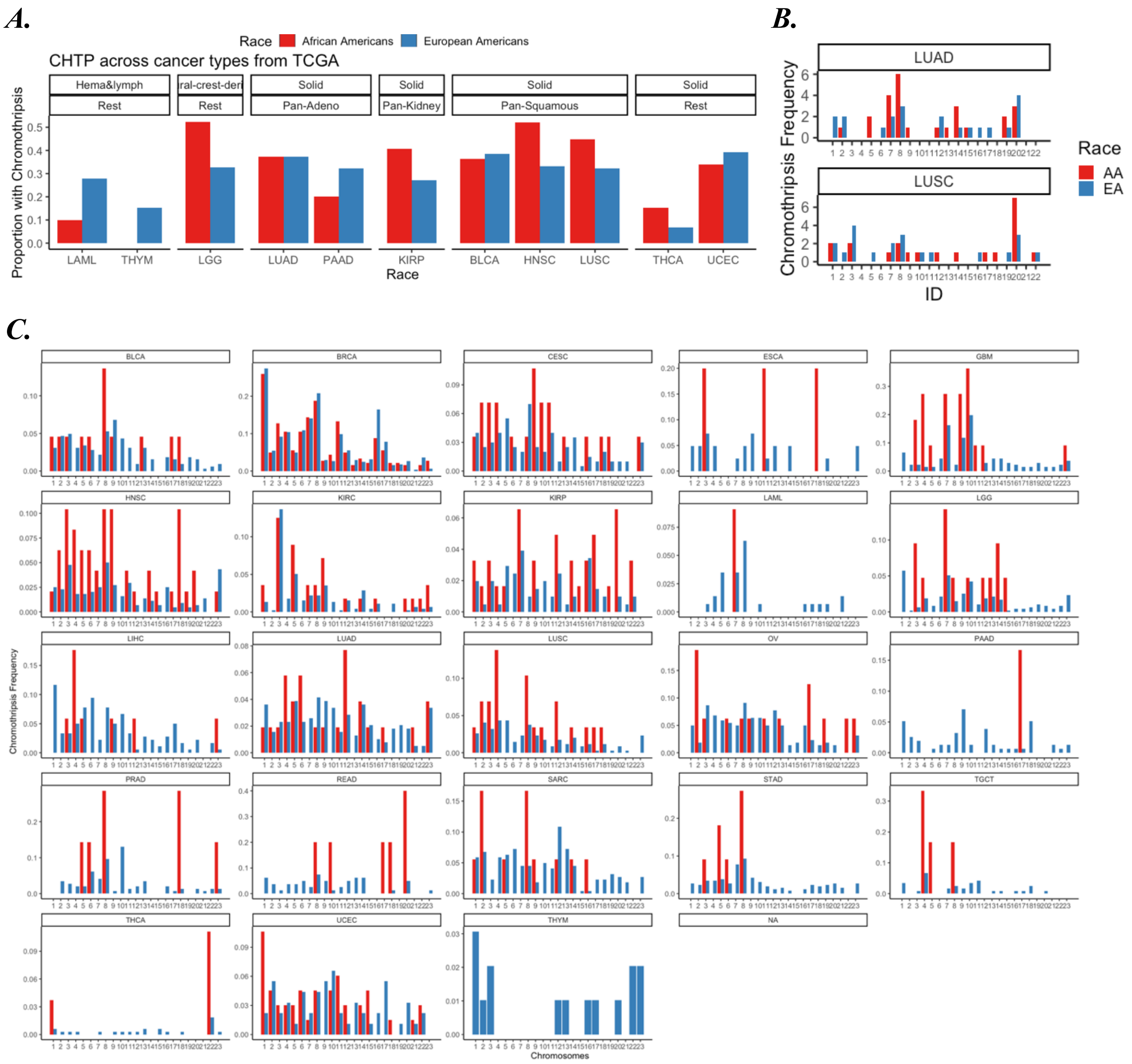


***Figure S7: A) Chromothripsis (CHTP) across race and chromosomes in TCGA and NCI-MD:***

***A)*** *Charting CHTP frequency distribution in AAs and EAs in various cancer types across TCGA.*

***B)*** *CHTP frequency across chromosomes for NCI-MD cohort in LUSC and LUAD C) CHTP frequency across chromosomes for various cancer type in TCGA cohort.*

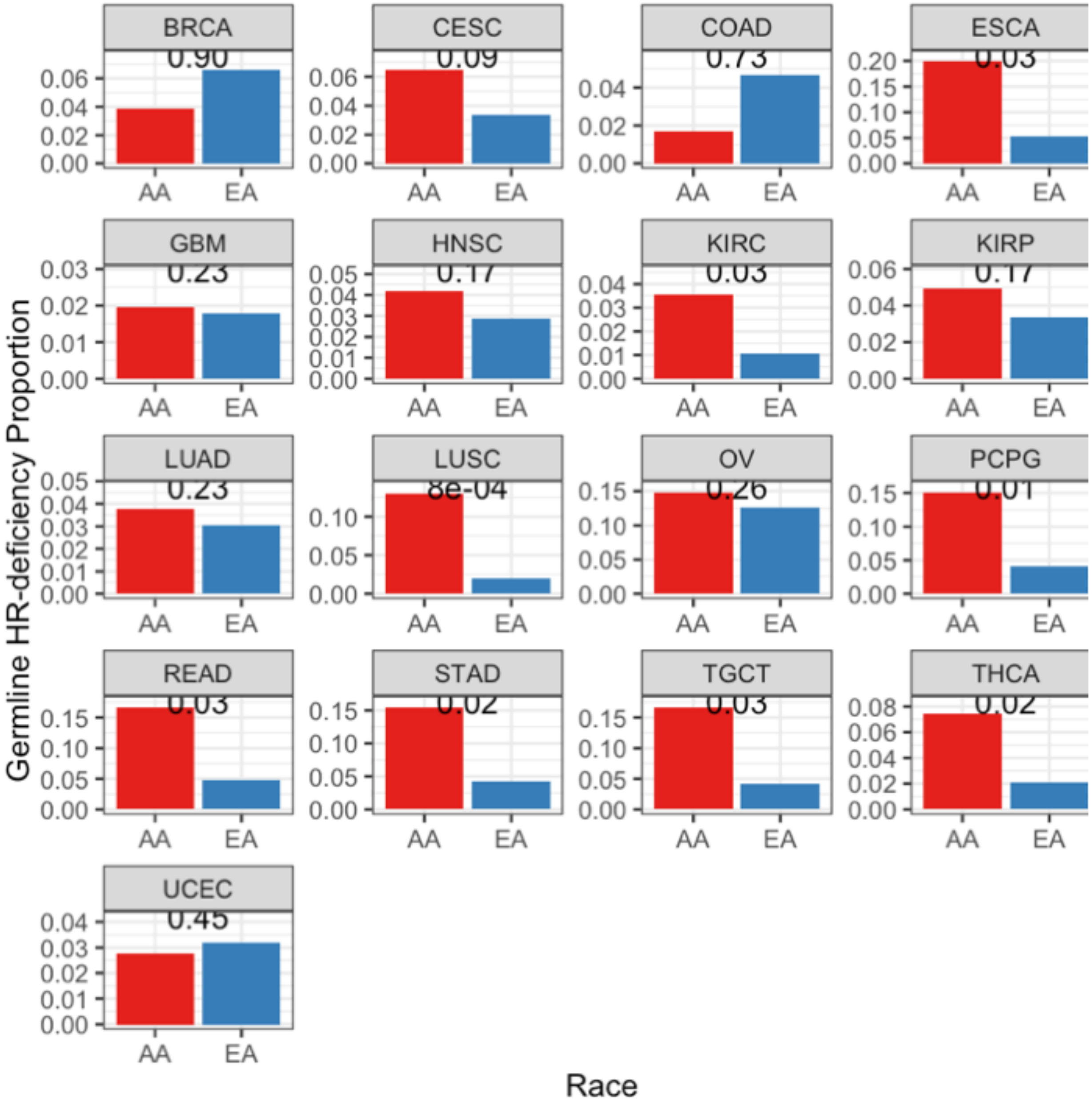


***Figure S8: Charting prevalence of germline predisposition*** *for HRD for 11 cancer types with at least 30 AA samples in TCGA, where HRD is defined by 108 hallmark genes.*

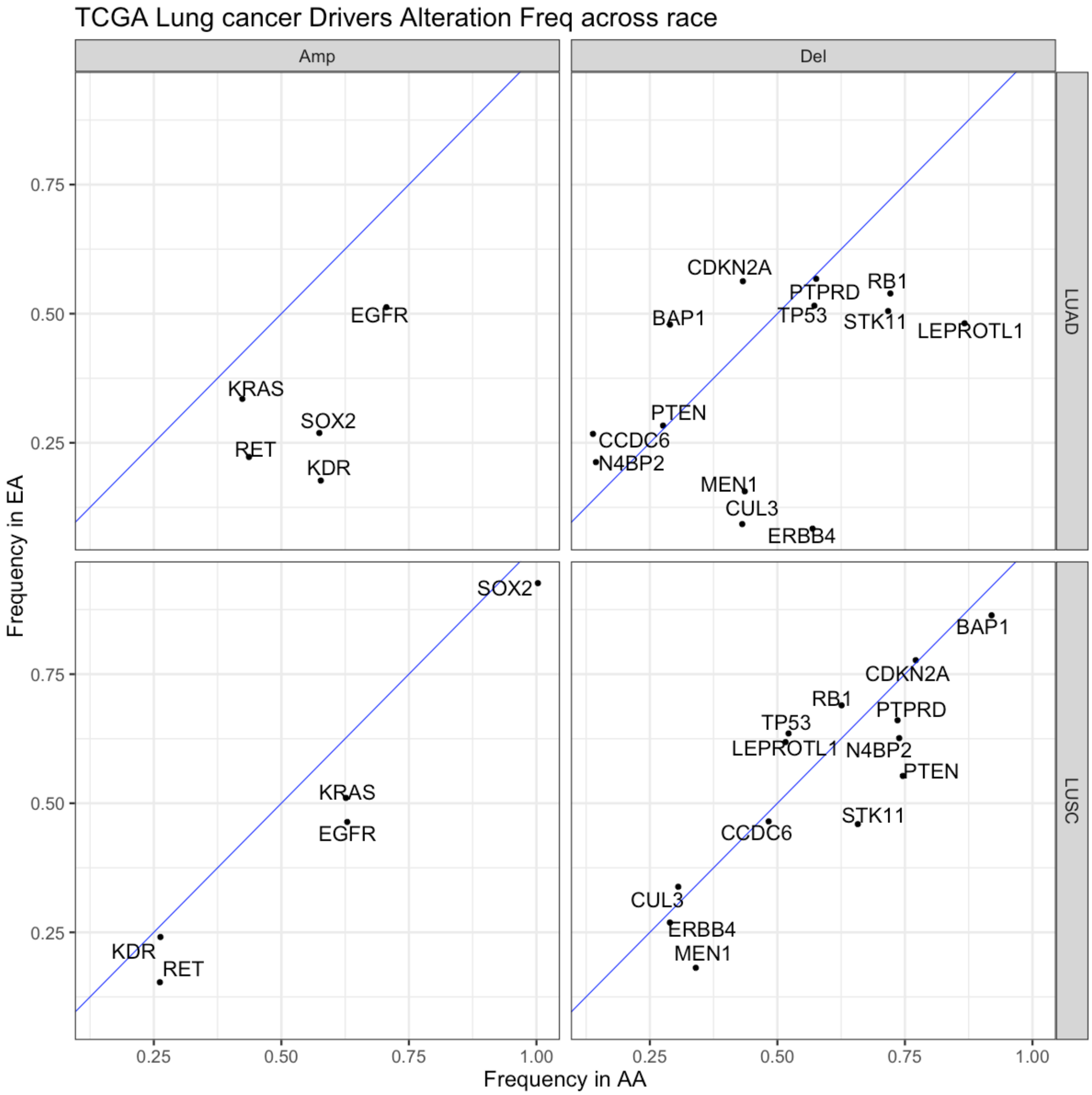


***Figure S9: SCNA of lung cancer drivers in LUSC and LUAD from TCGA:*** *Frequencies in EAs and AAs tumors in LUAD and LUSC from TCGA were plotted with blue diagonal line as neutral axis.*

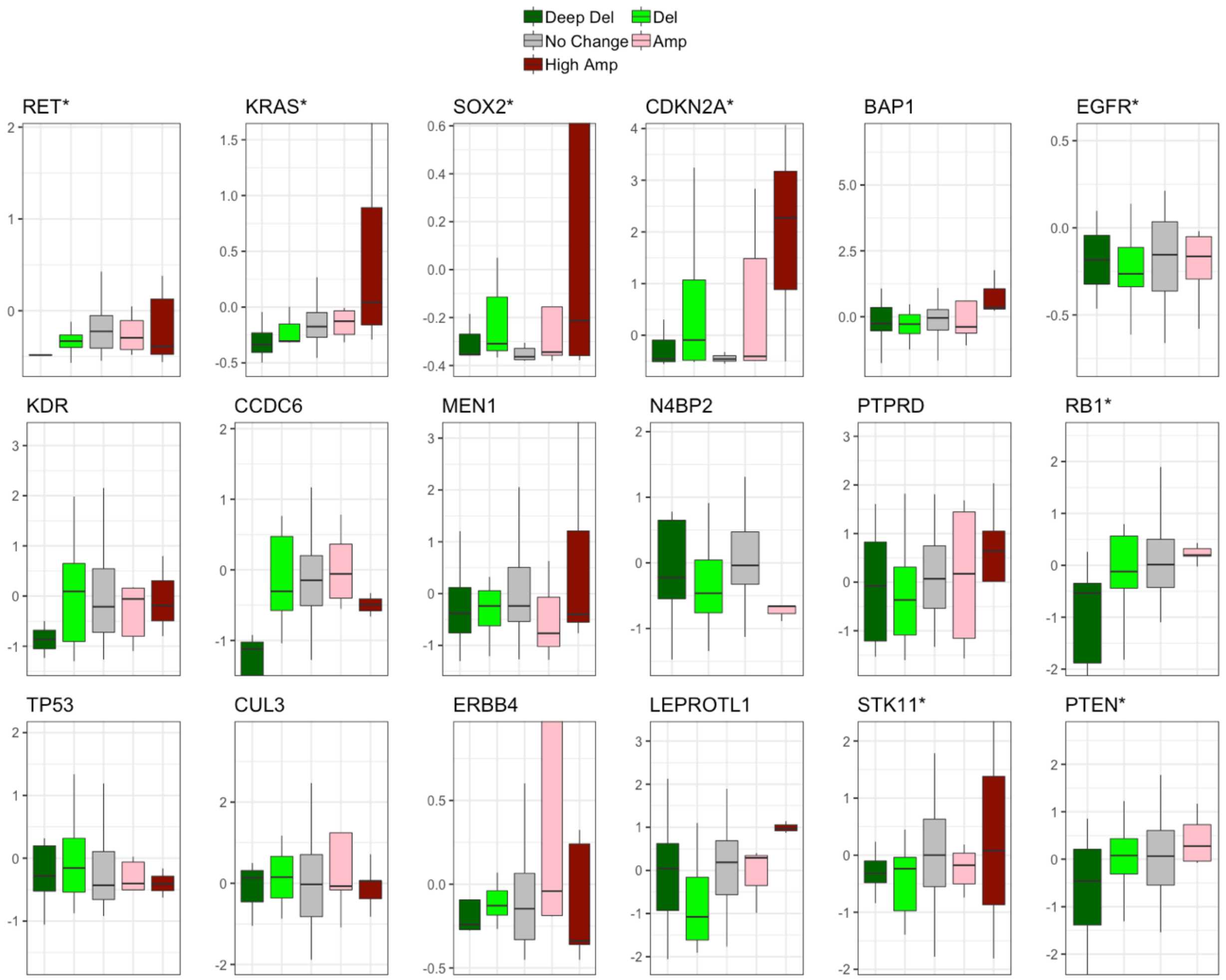


***Figure S10: Effect of copy number changes on expression*** *has been plotted for lung cancer drivers whose frequency across populations are significantly different. Asterisks alongside a gene name signifies a significant correlation between expression and CNV with a Spearman Rho>0.5.*

***References***


1. Hunt, Sarah E., et al. "Ensembl variation resources." Database 2018 (2018).
2. Forbes, Simon A., et al. "COSMIC: somatic cancer genetics at high-resolution." *Nucleic acids research* 45.D1 (2016): D777-D783.
3. Watkins, Johnathan A., et al. "Genomic scars as biomarkers of homologous recombination deficiency and drug response in breast and ovarian cancers." *Breast Cancer Research* 16.3 (2014): 211.
